# Comparative analyses of Gram-negative bacteria isolated from cancer patients with bacteraemia at the Uganda Cancer Institute

**DOI:** 10.64898/2026.07.05.736378

**Authors:** Margaret Lubwama, Lesley Hoyles, Anne L. McCartney, David P. Kateete, Freddie Bwanga, Edgar Kigozi, Leymon Kalema, Benon Asiimwe, George Katende, Fahad Lwigale, Simon Sekyanzi, Nixon Niyonzima, Jackson Orem, Henry Ddungu, Joyce Kambugu, Warren Phipps, Jody Winter

## Abstract

Antimicrobial resistance (AMR) exacerbates bacteraemia in cancer patients, particularly in low-resource settings. At the Uganda Cancer Institute, high rates of *Enterobacterales* producing extended-spectrum β-lactamases (ESBLs) have been reported, with DNA-based detection of *bla* genes limited to PCR. This study aimed to determine whether bacterial genomic DNA shipped at ambient temperature from Uganda to the UK retained sufficient quality for whole-genome sequencing (WGS), to allow in-depth genomic analyses of isolates.

Genomic DNA was extracted from Gram-negative bloodstream isolates (n=77) in Uganda and shipped to the UK at ambient temperature. *rpoB* gene (77/77, 100%) and WGS data (72/77, 93.5%) were generated for isolates, with 66/72 (91.7%) genomes of high-quality (*Escherichia coli* n=34; *Klebsiella* spp. n=32). Bioinformatic analyses included species identification, sequence typing, SNP analysis, AMR and virulence gene profiling, and comparison with publicly available genomes of Ugandan isolates.

Phenotypic–genotypic concordance was generally high: 7/77 (9.1%) isolates were misidentified by phenotypic testing, and two showed unexplained carbapenem resistance. *E. coli* isolates showed diverse sequence types, with high prevalence of *bla*_CTX-M_ (91.2%) and *bla*_OXA-1_ (47.1%); carbapenemase genes were rare. *Klebsiella* isolates lacked hypermucoidy loci and displayed diverse capsule types, with a high prevalence of ESBLs. Genomic clustering suggested limited within-hospital transmission of strains.

Genomic data can provide important insights into the dissemination of bacterial subclades of global concern. The widespread AMR genotypes reported here highlight the need for improved diagnostics and updated treatment guidelines for bacteraemia in Ugandan cancer patients.

**IMPACT STATEMENT:** Bloodstream infections are a major threat to cancer patients, particularly in low-resource settings where access to advanced diagnostics is limited and infection prevention may be challenging. This study shows that it is feasible to transport bacterial DNA at room temperature from Uganda to the UK for high-quality whole-genome sequencing, helping to overcome a logistical barrier to genomic surveillance.

By applying genomic analysis to bacteria that had caused bloodstream infections at the Uganda Cancer Institute, we found that traditional laboratory methods can misidentify some bacteria, and that genomic analyses can provide more accurate and detailed insights into the bacteria causing these serious infections.

Our work also highlights gaps in current treatment guidelines and demonstrates how genomic data could help inform updates to these, facilitating more effective antibiotic use in situations where urgent treatment is needed and there is no time to wait for laboratory test results.

While there was limited evidence of direct transmission of bacteria between patients in our dataset, the genetic diversity and resistance patterns we observed are concerning and emphasise the need for ongoing monitoring and more extensive future studies.

## INTRODUCTION

Bacteraemia in cancer patients is worsened by the burden of antimicrobial resistance (AMR), a growing public health concern associated with increased hospitalization and mortality. Globally, 4·71 million deaths were linked to bacterial AMR in 2021, with 1·14 million deaths caused by bacterial AMR. By 2050, it is estimated that 10.1 million deaths could occur (1). Hospitalized cancer patients have up to two times higher AMR rates compared to non-cancer patients, and the mortality rate of fatal infections in cancer patients is approximately three times that of the general population (2).

Multidrug-resistant (MDR) *Enterobacterales* are a significant cause of bacteraemia in cancer patients with increasing rates of third-generation cephalosporin-resistant *Enterobacterales* (3GCRE) and carbapenem-resistant *Enterobacterales* (CRE) reported. 3GCRE and CRE are both listed as critical priorities in the WHO bacterial priority pathogens list (3). A multicentre study in the US reported that extended-spectrum-beta-lactamase-producing *Enterobacterales* (ESBL-PE) were more likely to be isolated from patients with cancer versus patients without cancer (1).

Limited data are available for bacteraemia cases caused by MDR *Enterobacterales* across sub-Saharan Africa. Studies in Sudan and Zimbabwe reported ESBL-PE rates of 34.1% and 66.7%, respectively (4,5). In Uganda, our previous studies at the Uganda Cancer Institute (UCI), which focused on bacteraemia in hematologic cancer patients with febrile neutropenia, reported ESBL-PE phenotype rates of 74% (54/73) and genotype rates of 75% (55/73), with CTX-M the most prevalent ESBL (n=50/55, 91%). Genomic characterization of *Enterobacterales* isolated at the UCI was limited to ESBL genes (*bla*_CTX-M_, *bla*_TEM_, *bla*_SHV_) detected using PCR (6).

To fully characterize the *Enterobacterales* previously recovered from cancer patients at the UCI (6), here we used whole-genome sequencing (WGS) and bioinformatics analyses to generate extensive genotypic data for the clinical isolates. We compared our genotypic and phenotypic data to determine whether there was discordance between the two approaches (7,8). We also compared our newly generated sequence data with genomic data previously generated for *Enterobacterales* from Uganda and available from public repositories.

## METHODS

### Ethical approval and phenotypic characterization of isolates

Details of ethical approval and informed consent for the collection of isolates included in this study have already been reported (6). Previously phenotyped and stored *Enterobacterales* isolates were resuscitated from frozen (-80 °C) stocks and included in the current study (**Table 1**). Antimicrobial susceptibility tests were carried out using the Kirby Bauer disc diffusion method. Zone diameters of inhibition were interpreted according to the Clinical & Laboratory Standards Institute (CLSI) guidelines.

**Table 1.** Summary information for the genomic data generated in this study.

| Isolate | Date isolated | *rpoB* ID * | Coverage (x) | Completeness (%) | Contamination (%) | N50 | Contigs | Length (nt) | sourmash | ANI (%) # |
| --- | --- | --- | --- | --- | --- | --- | --- | --- | --- | --- |
| M1 \$ | 5/6/2019 | Kpn | 39 | 100 | 0.39 | 406280 | 34 | 5354870 | Kpn | Kpn 99.0% |
| M2 | 15/7/2019 | Eco | 53 | 100 | 0.02 | 214415 | 55 | 4728414 | Eco | Eco 96.7% |
| M3 | 10/12/2021 | Kpn | 39 | 100 | 0.23 | 138347 | 120 | 5310521 | Kpn | Kpn 98.9% |
| M4 | 7/12/2018 | Eco | 29 | 100 | 1.33 | 146848 | 161 | 5160174 | Eco | Eco 98.7% |
| M5 | 7/10/2021 | Eco | - | - | - | - | - | - | - | - |
| M6 | 16/11/2020 | Kox | 37 | 100 | 1.6 | 29692 | 512 | 6284704 | Kox | Kox 98.9% |
| M7 | 6/4/2019 | Eco | 42 | 100 | 1.11 | 250466 | 98 | 5348668 | Eco | Eco 98.2% |
| M8 | 27/3/2019 | Kpn | 20 | 100 | 0.33 | 25610 | 397 | 5145258 | Kpn | Kpn 98.8% |
| M9 | 22/5/2019 | Eco | 37 | 100 | 0.47 | 130539 | 105 | 4946726 | Eco | Eco 96.6% |
| M10 | 10/7/2019 | Eco | 40 | 100 | 0.02 | 183583 | 59 | 4727622 | Eco | Eco 96.7% |
| M11 | 10/8/2021 | Eco | 36 | 100 | 0 | 82858 | 217 | 4915801 | Eco | Eco 96.6% |
| M12 | 10/12/2020 | Eco | 73 | 100 | 0 | 309855 | 43 | 4898519 | Eco | Eco 96.8% |
| M13 | 27/3/2019 | Eco | 48 | 100 | 1.21 | 196250 | 92 | 5343195 | Eco | Eco 98.2% |
| M14 | 4/2/2019 | Kpn | 35 | 100 | 0.82 | 295707 | 65 | 5632375 | Kpn | Kpn 98.8% |
| M15 | 4/4/2019 | Kpn | 86 | 100 | 0.28 | 473648 | 54 | 5591184 | Kpn | Kpn 98.9% |
| M16 | 23/8/2018 | Eco | 31 | 100 | 0.33 | 162991 | 101 | 5271310 | Eco | Eco 96.7% |
| M17 | 10/10/2018 | Eco | 36 | 100 | 0.15 | 136264 | 87 | 4847410 | Eco | Eco 96.7% |
| M18 | 10/11/2020 | Kpn | - | - | - | - | - | - | - | - |
| M19 | 13/11/2018 | Kpn | 65 | 100 | 0.04 | 436237 | 40 | 5506850 | Kpn | Kpn 98.9% |
| M20 | 2/4/2019 | Eco | 37 | 100 | 0.74 | 136837 | 358 | 5367591 | Eco | Eco 96.6% |
| M21 | 30/5/2019 | Eco | 78 | 100 | 0.47 | 221145 | 63 | 5060625 | Eco | Eco 96.7% |
| M22 | 27/1/2020 | Kpn | 39 | 100 | 0.81 | 83471 | 136 | 5578248 | Kpn | Kpn 99.0% |
| M23 | 12/9/2018 | Eco | 86 | 100 | 0.7 | 329923 | 78 | 5184128 | Eco | Eco 98.1% |
| M24 | 4/5/2019 | Kpn | 30 | 100 | 0.12 | 430874 | 43 | 5419591 | Kpn | Kpn 98.9% |
| M25 | 16/11/2020 | Kpn | 6 | 27.32 | 0.18 | 894 | 1886 | 1671616 | - | - |
| M26 | 29/5/2018 | Kpn | 32 | 100 | 0.35 | 276866 | 78 | 5592070 | Kpn | Kpn 98.9% |
| M27 | 20/12/2017 | Kpn | 59 | 100 | 0.99 | 509719 | 62 | 5713335 | Kpn | Kpn 98.9% |
| M28 | 10/7/2019 | Eco | 66 | 100 | 0.02 | 189894 | 61 | 4727173 | Eco | Eco 96.7% |
| M29 | 31/8/2018 | Kpn | 43 | 100 | 0.2 | 295913 | 43 | 5535775 | Kpn | Kpn 99.0% |
| M30 | 4/2/2019 | Eco | 30 | 100 | 0.09 | 134984 | 99 | 4953041 | Eco | Eco 96.7% |
| M31 | 26/3/2019 | Eco | 32 | 100 | 0.16 | 142404 | 89 | 4851445 | Eco | Eco 96.7% |
| M32 | 20/3/2019 | Eco | 34 | 100 | 0.13 | 131189 | 105 | 4944018 | Eco | Eco 96.7% |
| M33 | 15/7/2019 | Eco | 47 | 100 | 0.19 | 139851 | 94 | 4857620 | Eco | Eco 96.6% |
| M34 | 14/2/2019 | Eco | 34 | 100 | 0.08 | 209705 | 54 | 4765818 | Eco | Eco 96.8% |
| M35 | 16/9/2021 | Kpn | 36 | 100 | 0.22 | 140179 | 109 | 5296247 | Kpn | Kpn 98.9% |
| M37 | 17/6/2018 | Kpn | 35 | 100 | 0.04 | 306769 | 57 | 5609154 | Kpn | Kpn 98.9% |
| M38 | 13/2/2019 | Kpn | 92 | 100 | 0.47 | 296151 | 41 | 5520673 | Kpn | Kpn 98.9% |
| M39 | 6/3/2018 | Kpn | 67 | 100 | 0.54 | 240543 | 42 | 5582814 | Kpn | Kpn 98.8% |
| M40 | 3/9/2018 | Eco | 103 | 100 | 0.13 | 165610 | 96 | 4946108 | Eco | Eco 96.7% |
| M41 | 15/2/2019 | Eco | 30 | 100 | 0.04 | 138727 | 102 | 4759446 | Eco | Eco 96.7% |
| M42 | 15/10/2018 | Eco | 42 | 100 | 0.28 | 210064 | 75 | 5241823 | Eco | Eco 97.0% |
| M43 | 23/5/2019 | Kqu | - | - | - | - | - | - | - | - |
| M44 | 23/5/2019 | Eho | 16 | 5.32 | 0 | 767 | 72 | 62292 | - | - |

| Isolate | Date isolated | rpoB ID * | Coverage (x) | Completeness (%) | Contamination (%) | N50 | Contigs | Length (nt) | sourmash | ANI (%) # |
| --- | --- | --- | --- | --- | --- | --- | --- | --- | --- | --- |
| M47 | 18/7/2018 | Eco | 41 | 100 | 0.41 | 185857 | 84 | 5251169 | Eco | Eco 96.8% |
| M49 | 11/5/2019 | Kpn | 42 | 100 | 0.14 | 115786 | 116 | 5535286 | Kpn | Kpn 99.0% |
| M55 | 16/11/2021 | Eco | 50 | 100 | 0.45 | 139897 | 100 | 5044137 | Eco | Eco 96.6% |
| M64 | 10/11/2020 | Eco | 49 | 100 | 0.1 | 242885 | 98 | 5065282 | Eco | Eco 96.7% |
| M67 | 28/10/2020 | Eco | 60 | 100 | 0.16 | 126451 | 129 | 4926442 | Eco | Eco 96.7% |
| M68 | 29/11/2021 | Eco | 47 | 100 | 0.85 | 294772 | 85 | 5234839 | Eco | Eco 98.6% |
| M70 | 17/12/2021 | Eco | 35 | 100 | 1.42 | 229877 | 74 | 5146136 | Eco | Eco 98.3% |
| M82 | 2/4/2019 | Eco | 4 | 6.3 | 0 | 4004 | 4 | 5799 | - | - |
| M83 | 2/4/2019 | Kpn | 49 | 100 | 1.82 | 193098 | 105 | 5823584 | Kpn | Kpn 98.9% |
| M84 | 8/10/2021 | Kpn | 34 | 100 | 0.21 | 353958 | 83 | 5812225 | Kpn | Kpn 98.9% |
| M88 | 27/6/2018 | Kpn | 53 | 100 | 0.26 | 338569 | 49 | 5646519 | Kpn | Kpn 98.9% |
| M89 | 7/10/2019 | Eco | 40 | 100 | 0.21 | 172992 | 82 | 5109503 | Eco | Eco 96.8% |
| M90 | 20/1/2021 | Kqu | 34 | 100 | 0.44 | 225655 | 69 | 5468044 | Kqu | Kqu 96.4% |
| M91 | 20/1/2021 | Kqu | 52 | 100 | 0.34 | 184970 | 76 | 5479188 | Kqu | Kqu 96.4% |
| M92 | 30/9/2019 | Kva | 38 | 100 | 0.35 | 469657 | 28 | 5507202 | Kva | Kva 96.9% |
| M93 | 27/9/2021 | Eco | 54 | 100 | 0.23 | 178075 | 83 | 5196754 | Eco | Eco 96.8% |
| M94 | 29/11/2021 | Kpn | 54 | 100 | 0.29 | 285643 | 48 | 5432432 | Kpn | Kpn 99.0% |
| M95 | 10/9/2019 | Kpn | 44 | 100 | 0.19 | 213291 | 78 | 5679219 | Kpn | Kpn 98.9% |
| M96 ^ | 11/7/2019 | Kpn | 36 | 100 | 0.52 | 442094 | 68 | 5675302 | Kpn | Kpn 98.9% |
| M97 | 22/9/2018 | Eco | 65 | 100 | 0.18 | 134895 | 80 | 4810923 | Eco | Eco 96.7% |
| M98 | 2/9/2019 | Eco | 34 | 6.38 | 0 | 4754 | 5 | 18981 | - | - |
| M99 | 23/5/2019 | Kpn | 53 | 100 | 0.12 | 185943 | 95 | 5317221 | Kpn | Kpn 99.0% |
| M100 | 5/10/2020 | Kpn | 51 | 100 | 0.15 | 370220 | 80 | 5844935 | Kpn | Kpn 98.8% |
| M101 | 20/2/2019 | Eco | 56 | 100 | 0.78 | 284400 | 47 | 5022629 | Eco | Eco 98.6% |
| M102 | 16/12/2019 | Eco | 12 | 81.44 | 4.2 | 2076 | 2496 | 3859008 | - | - |
| M103 | 1/10/2018 | Eco | 47 | 100 | 0.72 | 270599 | 54 | 5202972 | Eco | Eco 98.2% |
| M104 | 15/11/2019 | Eco | 61 | 100 | 0.34 | 139580 | 98 | 4888437 | Eco | Eco 96.6% |
| M105 | 2/4/2019 | Kpn | - | - | - | - | - | - | - | - |
| M106 | 16/12/2019 | Kpn | 126 | 100 | 0.84 | 287874 | 52 | 5424042 | Kpn | Kpn 98.9% |
| M107 | 17/1/2019 | Eco | - | - | - | - | - | - | - | - |
| M108 ^ | 11/7/2019 | Kpn | 98 | 100 | 0.49 | 442094 | 59 | 5679703 | Kpn | Kpn 98.9% |
| M109 | 23/5/2019 | Eho | 5 | 8.55 | 0.01 | 734 | 410 | 320776 | - | - |
| M110 \$ | 5/6/2019 | Kpn | 92 | 100 | 0.4 | 464663 | 23 | 5355503 | Kpn | Kpn 99.0% |
| M111 | 20/1/2021 | Kpn | 79 | 100 | 0.76 | 421071 | 56 | 5508552 | Kpn | Kpn 98.9% |

\* Eco, *Escherichia coli*; Eho, *Enterobacter hormaechei*; Kox, *K. oxytoca*; Kpn, *K. pneumoniae*; Kqu, *K. quasipneumoniae* subsp. *quasipneumoniae*; Kva, *K. variicola*.

### DNA extraction

DNA was extracted from isolates using the modified cetyltrimethylammonium bromide method (figshare doi:10.6084/m9.figshare.32688180). DNA was quantified and its purity checked ( *A*_260_/*A*_280_ ratio) using a Nanodrop spectrophotometer. For long-term storage, DNA was kept at -80 °C in cryovials.

DNA suspended in nuclease-free water was transported via FEDEX to the microbiology laboratory of Nottingham Trent University, UK at ambient temperature (shipping time ∼3 days).

### Quality control of shipped DNA

Upon receipt in the UK, DNA samples were stored at 4 °C (1–8 weeks). DNA was quantified using a Qubit 4 Fluorometer (1✕ dsDNA HS kit; Invitrogen). DNA fragmentation was assessed on a 4200 TapeStation system (Agilent).

### *rpoB* gene-based sequence analysis

*rpoB* genes were PCR-amplified from DNA (9), with PCR products cleaned (NucleoSpin Gel and PCR Clean-up kit; Marchery Nagel) and Sanger-sequenced (Source BioScience; Nottingham, UK). Sequence data were processed as recommended (9) (figshare doi:10.6084/m9.figshare.31534144), and compared with publicly available *rpoB* gene sequences using NCBI BLASNT and aligned against publicly available data using CLUSTAL W (Geneious Prime v2025.2.1).

### WGS and bioinformatics

Samples (DNA >1 ng/µl) were sent to MicrobesNG (Birmingham, UK) for Illumina short-read sequencing and genome assembly (10). Returned assembled genomes were filtered using BBmap: contigs >500 nt in length were retained. Completeness and contamination of assemblies were assessed using CheckM2 v1.1.0 (11). sourmash v4.9.4 (lca, GTDB rs214) was used to identify isolates based on genome sequence data (12). fastANI v1.3.4 (13) was used to compare genome sequences with those of type strains of closest relatives. Genes encoded in genomes were predicted and annotated (figshare doi:10.6084/m9.figshare.32688555) using Bakta (software v1.11.3; database v6.0; AMRFinder v4.0.23, database 2025-07-16.1) (14).

Kaptive v3.1.0 was used to type K and O loci of *Klebsiella* spp. (15). Kleborate v3.2.4 (AMRFinder v4.0.23, database v2025-07-16.1) was used to characterize *E. coli* isolates and members of the *K. pneumoniae* species complex (KpSC) and *K. oxytoca* species complex (KoSC) (16). MLST data generated for KoSC (PubMLST), *E. coli* (PubMLST) and KpSC (BIGSdb-Pasteur) were uploaded to the indicated public MLST databases. antiSMASH v8.0.4 (17) and data included elsewhere (18,19) were used to identify biosynthetic gene clusters (BGCs) [tilimycin (*til*) and leupeptin (*leup*), respectively] encoded by members of the KoSC (figshare doi:10.6084/m9.figshare.31546765, doi:10.6084/m9.figshare.31545100, doi:10.6084/m9.figshare.31541713). *E. coli* sequence data were uploaded to EnteroBase v1.2.0 (20) on 25 November 2025, for comparison with other genomic data from Uganda and SNP-based analysis. Roary v3.13.0 (21) and SNP-sites v2.5.1 (input core gene alignment file from Roary) (22) were used to analyse KpSC genome data. Additional published genomic data for *E. coli* (23–28) and relevant *Klebsiella* spp. (23–25,29–33) from Uganda were included in comparative analyses. MLST data for 192 Ugandan KpSC isolates were downloaded from BIGSdb-Pasteur.

## RESULTS

Seventy-seven of 83 *Enterobacterales* isolates recovered from the blood of haematologic cancer patients with febrile neutropenia in an adult ward and a paediatric ward at the UCI between November 2017 and December 2021 could be resuscitated (**Table 1**). We aimed to determine whether we could send high-quality DNA to the UK at ambient temperatures by post for WGS, to use genomic data generated to further characterize UCI isolates, and to compare the AMR genotypes of these isolates with their previously reported phenotypic characteristics (6). On arrival in the UK, DNA was of sufficient quantity and integrity for sequencing (58.5 +/- 32.1 ng/µl, n=77; DNA Integrity Number 8.4 +/- 0.8, n=49; **Supplementary Table 1).**

### *rpoB* gene-based identification

Initially *rpoB* gene sequence analysis was used to identify the 77 isolates, as this approach is more effective for species-level identification of Gram-negative bacteria than 16S rRNA gene sequence-based analysis (34). From most to least prevalent, the UCI isolates represented *Escherichia coli* (n=39, 50.6%), *Klebsiella pneumoniae* (n=31, 40.3%), *Klebsiella quasipneumoniae* (n=3, 3.9%), *Enterobacter hormaechei* (n=2, 2.6%), *Klebsiella oxytoca* (n=1, 1.3%) and *Klebsiella variicola* (n=1, 1.3%) (**Table 1**). Discordance between phenotypic and genotypic identification was seen in 7/77 (9.1%) isolates: *K. oxytoca* (n=1) was identified as *E. coli* in (6); *K. quasipneumoniae* (n=2) and *K. variicola* (n=1) were identified as *K. pneumoniae* in (6); three isolates originally identified as *Enterobacter* spp. (6) were found to be *K. quasipneumoniae* (n=1) and *Enterobacter hormaechei* (n=2) in this study.

### Genome-based identification of isolates

Genome data could be generated for 72/77 isolates: 66 represented high-quality (35) draft genome sequences that were included in further analyses (**Table 1**). Insufficient DNA (in terms of volume and/or concentration) meant we could not generate high-quality sequence data for all isolates.

Species-level identification of the 66 isolates was confirmed using ANI- and sourmash-based analyses (36,37), with their findings in agreement and matching *rpoB* gene-based identification (**Table 1**). The high-quality genomes represented *E. coli* (n=34), *K. pneumoniae* (n=28), *K. quasipneumoniae* subsp. *similipneumoniae* (n=2), *K. variicola* subsp. *tropica* (n=1) and *K. oxytoca* (n=1).

### Escherichia coli

No UCI *E. coli* isolates (n=34) were predicted to encode for shiga toxin, *eae* (intimin), *ipaH* (ubiquitin ligase associated with enteroinvasive *E. coli*) or resistance to colistin by Kleborate. Enteropathogenic, shiga-toxin-encoding or *eae*-positive strains were rarely detected among publicly available Ugandan *E. coli* genomes [6/627 (1.0%), 15/627 (2.4%) and 6/627 (1.0%), respectively]; no *ipaH*-positive strains were detected, nor were any of the publicly available genomes predicted to encode genes for colistin resistance (**Supplementary Table 2**, **Supplementary Table 3**).

Sequence type (ST) ST167 was most abundant across UCI *E. coli* isolates (n=10 genomes), followed by ST131 (n=5), ST744 and ST405 (n=3 genomes each), ST38, ST410, ST617 and ST1193 (n=2 genomes each), and ST12, ST609, ST2083, ST361 and ST648 (n=1 genome each) (**Figure 1a**). Comparison of our MLST data with those for other *E. coli* (n=627) recovered in Uganda (**Supplementary Table 2**, **Supplementary Table 3**) showed UCI isolates included underrepresented STs for the country (**Figure 1a**). A total of 195 STs were represented across the entire Ugandan dataset: ST167 (62/661, 9.4%) > ST10 (60/661, 9.1%) > (36/661, 5.4%); all other STs represented <4% of the total.

**Figure 1.**
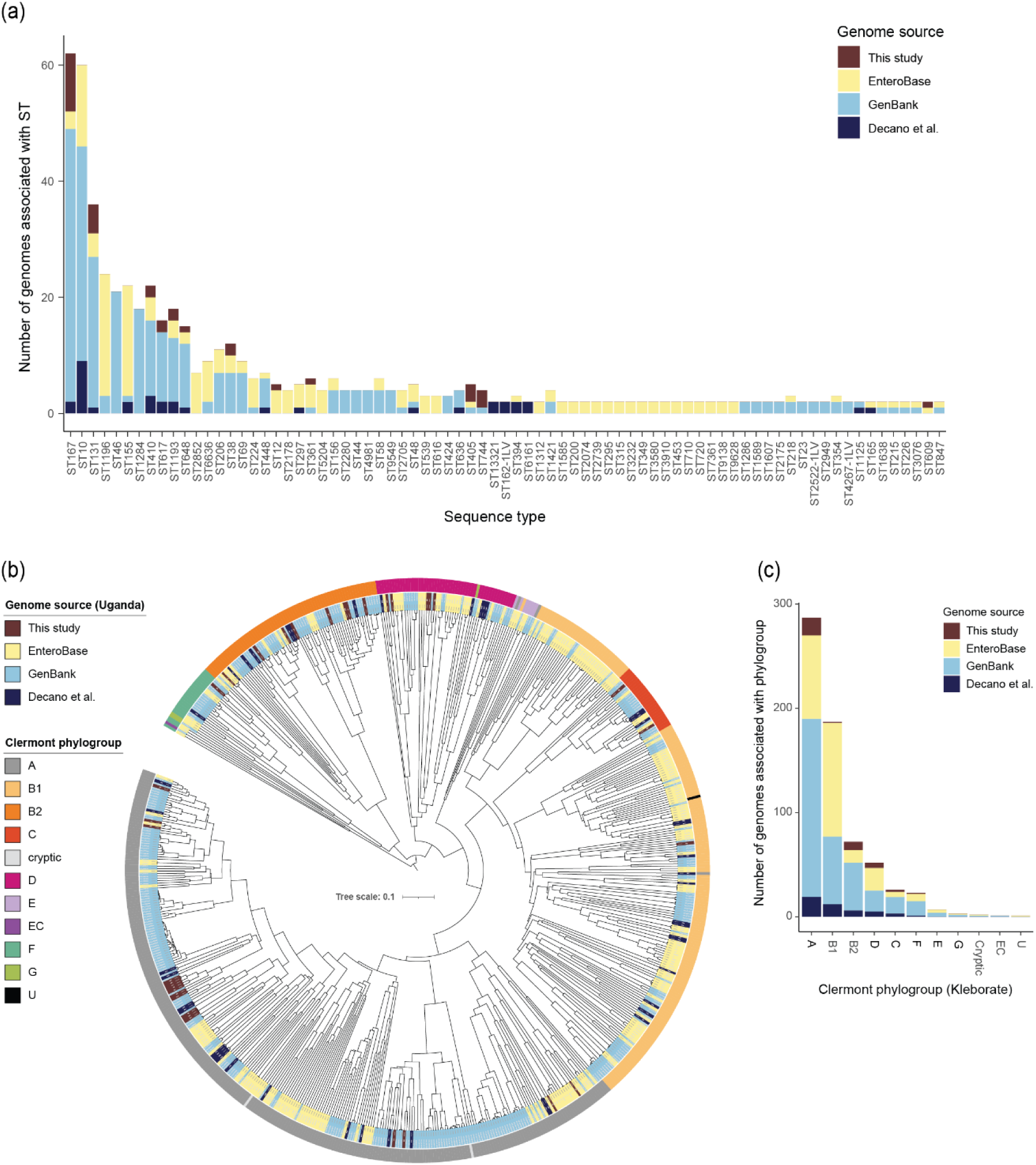
Comparison of genomic data derived from *E. coli* isolates recovered in Uganda. (a) Graphical summary of MLST data for 661 *E. coli* genomes [n=34, this study; n=241, EnteroBase; n=340, GenBank; n=46, outpatient urine isolates (25)]. STs represented only once in the dataset (n=117) were excluded from the figure, but are included in **Supplementary Tables 2** and **Supplementary Table 3**. (b) Dendrogram generated from sourmash kmer signatures showing similarity of the genomes from different sources and their Clermont phylogroup affiliations. The tree is rooted at the midpoint. Colours of the inner circle relate to the sources of the genomes, while those of the outer circle relate to the Clermont phylogroup affiliations; refer to colour legends shown in figure. (c) Bar graph summarizing Clermont phylogroup data for the 661 genomes included in the analysis.

We used sourmash (kmer signatures) to determine how similar UCI *E. coli* genomes were to others recovered from Uganda, and Kleborate to provide Clermont phylogroup information (**Figure 1b**, **1c**). Half (n=17/34) of our bloodstream isolates belonged to Clermont phylogroup A, followed by phylogroups B2 (n=8/34, 23.5%), D (n=5/34, 14.7%), C (n=2/34, 5.9%), B1 (n=1/34, 2.94%) and F (n=1/34, 2.9%) (**Figure 1c**). Most Ugandan genomes belonged to phylogroup A (287/661, 43.4%), followed by B1 (187/661, 28.3%), B2 (72/661, 10.9%), D (52/661, 7.9%), C (26/661, 3.9%), F (23/661, 3.5%) and E (7/661, 1.1%).

All UCI isolates were predicted to encode *bla*_TEM-1_ (33/34, 97.1%), except for M33 (*bla*_TEM-169_). Among the 627 public *E. coli* genomes several *bla*_TEM_ variants were found (**Supplementary Figure A**): *bla*_TEM-1_, 309/627 (49.3%); *bla*_TEM-135_, 10/627 (1.6%); *bla*_TEM-57_, 9/627 (1.4%); *bla*_TEM-30_, 5/627 (0.8%); *bla*_TEM-169_, 2/627 (0.3%).. *bla*_CTX-M-15_ dominated among UCI isolates (21/34, 61.8%), with *bla*_CTX-M-27_ (3/34, 8.8%) and *bla*_CTX-M-55_ (7/34, 20.6%) also detected; three isolates (3/34, 8.8%) were not predicted to encode a *bla*_CTX-M_ gene. *bla*_CTX-M-15_ dominated across the wider Ugandan *E. coli* genome dataset (309/627, 49.3%), followed by *bla*_CTX-M-27_ (21/627, 3.3%), *bla*_CTX-M-14_ (19/627, 3.0%), *bla*_CTX-M-55_ (7/627, 1.1%), *bla*_CTX-M-3_ (4/627, 0.6%), *bla*_CTX-M-24_ (2/627, 0.3%) and *bla*_CTX-M_, *bla*_CTX-M-137_ and *bla*_CTX-M-25_ (each 1/627, 0.2%). The *bla*_OXA-1_ gene was detected at high prevalence across UCI isolates (16/34, 47.1%), but was less prevalent in the Ugandan *E. coli* genome dataset (102/627, 16.3%).

Genomic data for some UCI isolates clustered together (**Figure 1b**), but the diversity of the entire Ugandan genome dataset required us to undertake a stand-alone EnteroBase analysis of our 34 genomes (**Figure 2**). Our genomes clustered by ST, and within some ST clusters there were pairs or groups of related strains. For example, the ST167 O8:H9 isolates M32 and M40 had identical AMR genotypes and phenotypes but differed by 64 SNPs (**Supplementary Table 4**). Within the ST131 cluster, isolates M7, M13, M70 and M103 were highly similar with only minor differences in AMR genotypes and phenotypes, but differed by 137-547 SNPs (**Supplementary Table 4**). Within the ST405 cluster, isolates M16 and M93 differed by 223 SNPs (**Supplementary Table 4**).

**Figure 2.**
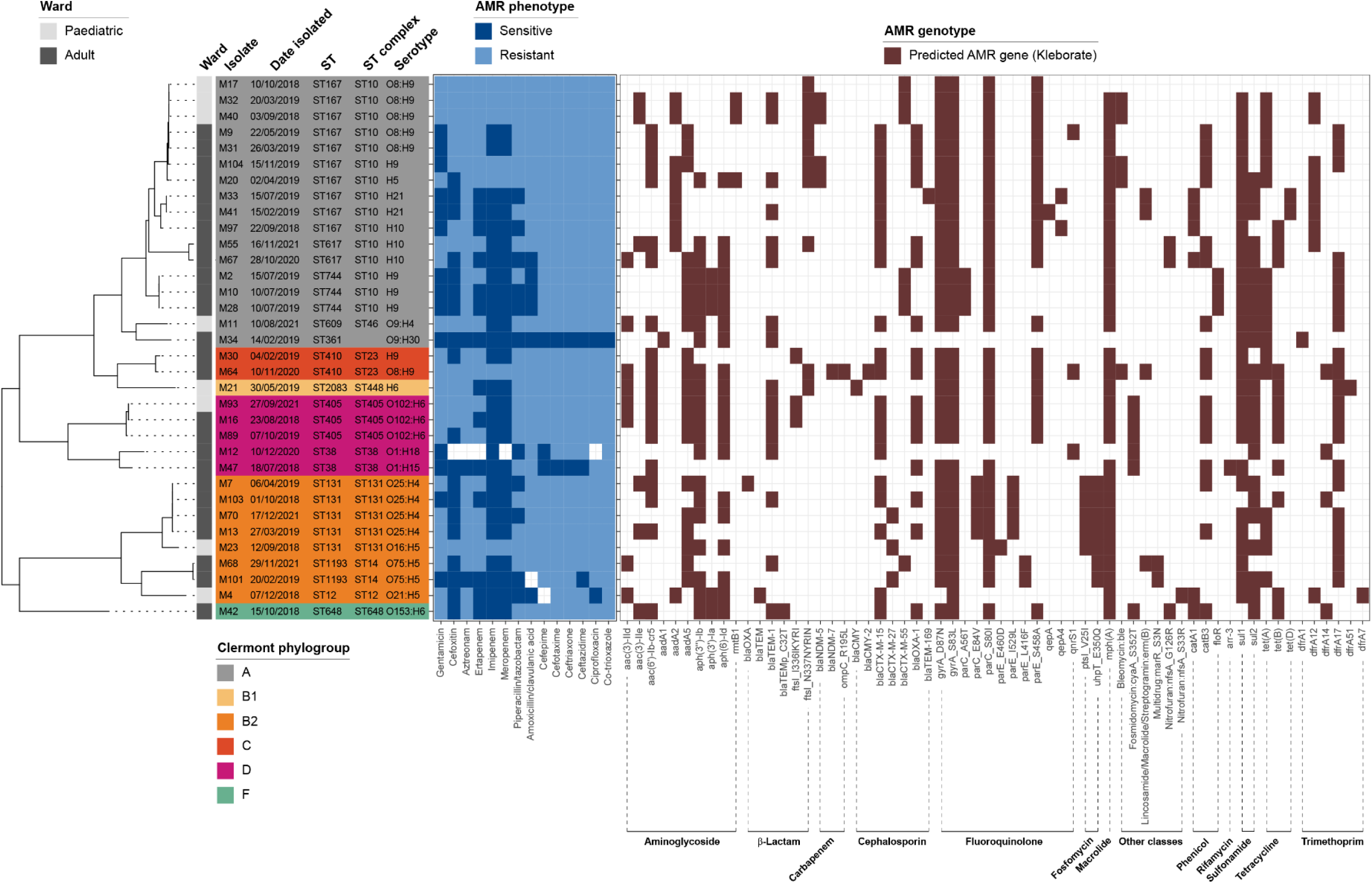
Dendrogram generated from EnteroBase analysis of our 34 *E. coli* genomes, along with ward sources of isolates, AMR phenotypic data and AMR gene predictions generated by Kleborate for the genomic data. The genome of isolate M2 was chosen at random as the reference sequence for the SNP-based analysis, which was run to detect a minimum of 95% sites present. Antibiotics used to generate phenotypic data: aminoglycoside – gentamicin; beta-lactam – cefoxitin; aztreonam (monobactam), ertapenem (carbapenem), imipenem (carbapenem), meropenem (carbapenem), piperacillin/tazobactam (ESBL), amoxicillin/clavulanic acid (penicillin), cefepime, cefotaxime, ceftriaxone and ceftazidime (all cephalosporins); fluoroquinolone – ciprofloxacin; sulfonamide – co-trioxazole.

AMR genotypes and phenotypes were broadly consistent, although phenotypic carbapenem resistance was observed in isolates M17 and M23 in the absence of *bla*_NDM_ or other carbapenemase-encoding genes. M34, the only pan-susceptible isolate, was also the only isolate without any predicted cephalosporin or fluoroquinolone resistance genes (**Figure 2**).

### Klebsiella pneumoniae species complex

High-quality genome sequences were generated for 31 UCI isolates belonging to the KpSC, which includes *K. pneumoniae*, *K. quasipneumoniae* and *K. variicola* (**Table 1**). Fourteen of 31 (45.2%) isolates were predicted to encode the virulence factor yersiniabactin (siderophore) (**Supplementary Table 5**). Across all Ugandan KpSC genomes included in this study (n=31, this study; n=106, publicly available (13,16,19–23)), none was predicted to encode the genotoxin colibactin; 72/137 (52.6%) encoded no siderophores, 63/137 (45.99%) encoded only yersiniabactin and 2/137 (1.5%) encoded only aerobactin (**Figure 3a**).

**Figure 3.**
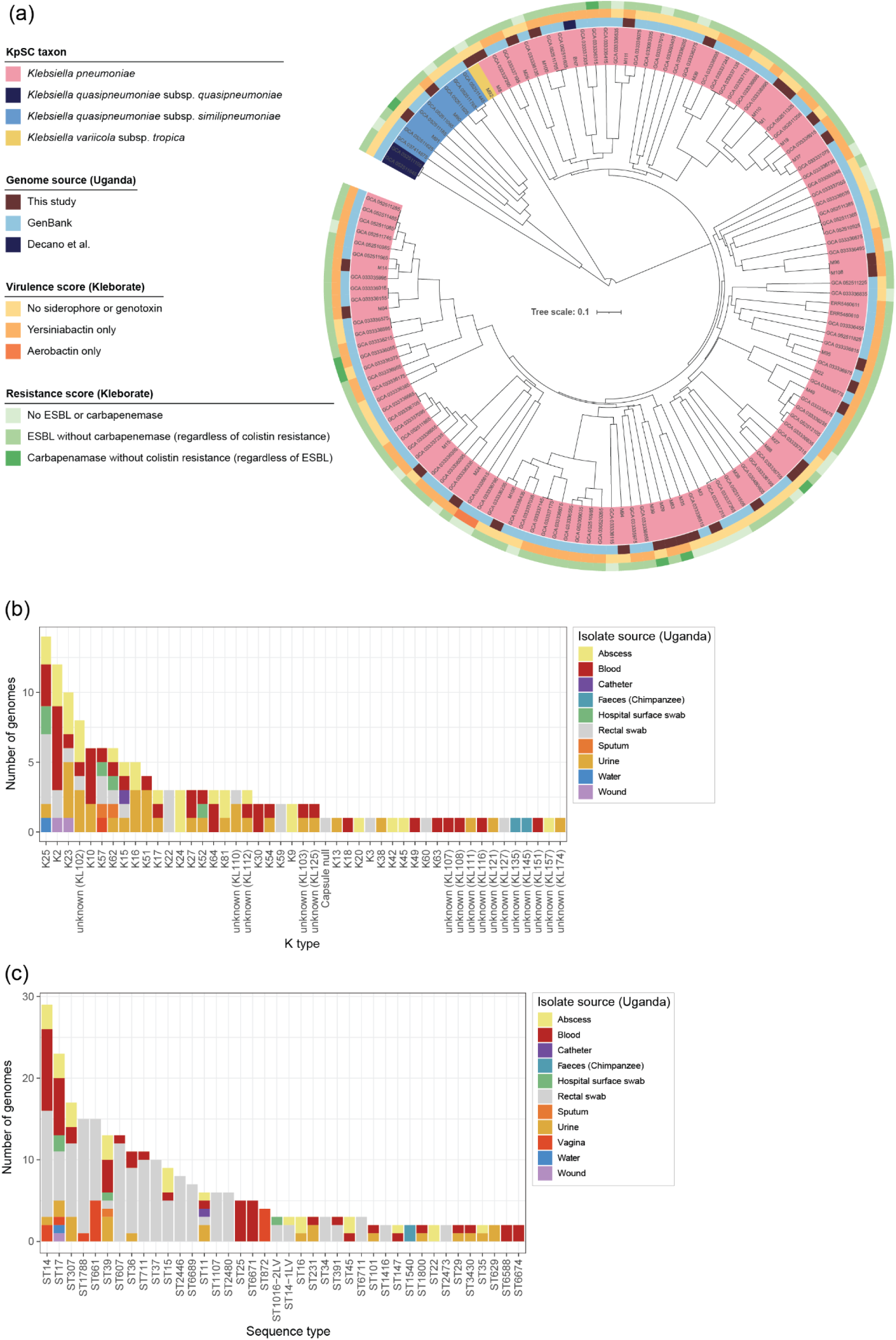
Comparison of genomic data derived from KpSC isolates recovered in Uganda. (a) Dendrogram generated from sourmash kmer signatures for 137 KpSC genomes [n=31, this study; n=136, GenBank; n=1, (25)]. The tree is rooted at the midpoint. (b) Bar graph summarizing Kleborate K (capsule) type data for the 137 genomes. (c) Bar graph summarizing MLST data for 329 KpSC isolates from Uganda (n=31, this study; n=106, public genome data; n=192 isolates, BIGSdb-Pasteur). All data are from human sources, with the exception of those for the water and chimpanzee isolates (**Supplementary Table 5**, **Supplementary Table 6**). Of the 106 different STs detected across the whole dataset, only those detected more than once are shown in the bar graph. (b, c) Bars are coloured based on source of isolation.

Kleborate identified 46 different K (capsule) types across the 137 Ugandan *Klebsiella* genomes (**Figure 3b**): K25 (14/137, 10.2%) > K2 (12/137, 8.8%) > K23 (10/137, 7.2%) > unknown KL102-like (8/137, 5.8%) > K10, K57 and K62 (all 6/137, 4.4%). One isolate (*K. pneumoniae* DKB0822) was reported as “capsule null”. K2 dominated the UCI isolates (6/31, 19.4%), followed by K10 and K25 (both 3/31, 9.7%), and K27, K30 and K64 (all 2/31, 6.5%). Eight O (LPS) types were represented in our genomes (O1αβ,2α – 11/31, 35.5%; O1αβ,2β – 8/31, 25.8%; O5 – 4/31, 12.9%; O2β – 3/31, 9.7%; O3αβ – 2/31, 6.5%; O14, O10, O13 – all 1/31, 3.2%) (**Figure 4a**). Eleven O types were found across all 137 genomes analysed here: O1αβ,2α (38/137, 27.7%) > O1αβ,2β (28/137, 20.4%) > O2β (15/137, 10.9%) > O5 (14/137, 10.2%) > O3αβ (8/137, 5.8%) (**Supplementary Table 5**). No UCI isolates were predicted to encode the capsule-associated *rmpADC* hypermucoidy locus, and only 2/137 (1.5%) of all Ugandan KpSC genomes could be assigned an RmST (16), namely *K. pneumoniae* MUWRP4500 (K2:O1αβ,2α) and MUWRP1041 (K57:O3γ).

**Figure 4.**
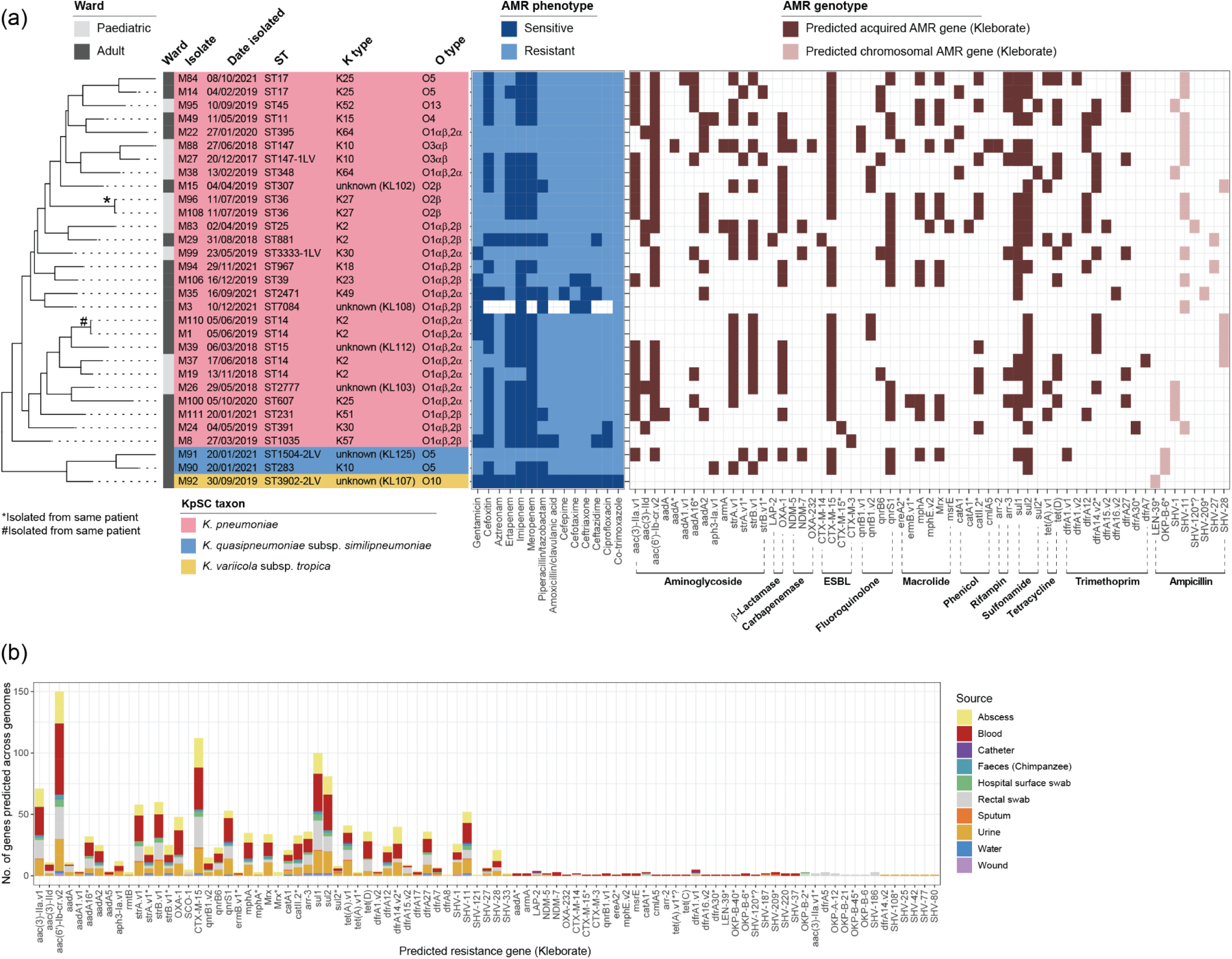
AMR profiles of KpSC isolates from Uganda. (a) Dendrogram generated from Roary-based analysis of our 31 KpSC genomes, along with ward sources of isolates, AMR phenotypic data and AMR gene predictions generated by Kleborate for the genomic data. Antibiotics used to generate phenotypic data: aminoglycoside – gentamicin; beta-lactam – cefoxitin; aztreonam (monobactam), ertapenem (carbapenem), imipenem (carbapenem), meropenem (carbapenem), piperacillin/tazobactam (ESBL), amoxicillin/clavulanic acid (penicillin), cefepime, cefotaxime, ceftriaxone and ceftazidime (all cephalosporins); fluoroquinolone – ciprofloxacin; sulfonamide – co-trioxazole. (b) Genotypic profiles of all KpSC genomic data available for Uganda (including our 31 genomes from blood): abscess, n=29; blood, n=38; catheter, n=1; faeces (Chimpanzee), n=2; hospital surface swab, n=5; rectal swab, n=23; sputum, n=2; urine, n=34; water, n=1; wound, n=2. (a, b) No distinction is made between sequences that have identical amino acid sequences but different nucleotide sequences compared with reference genomes, but full details are included in **Supplementary Table 5**. * suffix with gene names: no exact match at amino acid or nucleotide level found by Kleborate; the reported allele is the closest nucleotide match with the reference sequence.

MLST data were available for 329 Ugandan KpSC isolates (**Figure 3c**; **Supplementary Table 5**, **Supplementary Table 6**), representing 106 different STs. Twenty-six different STs were represented across our 31 genomes, with only ST14 (n=4), ST17 (n=2) and ST36 (n=2) represented more than once. The high-risk *K. pneumoniae* clones ST147 (isolate M88) and ST307 (isolate M15) (38) were represented among the 31 UCI isolates. An additional ST147 isolate, recovered from an abscess, was found among publicly available genome data, along with 16 ST307 isolates recovered from abscesses (n=3), blood (n=2), rectal swabs (n=9) and urine (n=3).

Carbapenemase- and ESBL-encoding genes were absent from 3/31 UCI isolates (9.7%; M3, M35, M92), with 27/137 (19.7%) of all Ugandan KpSC genomes lacking these genes (**Figure 3a**, **Figure 4a**). *K. pneumoniae* M3 was phenotypically susceptible to piperacillin-tazobactam, cefotaxime, ceftriaxone, imipenem and gentamicin; *K. pneumoniae* M35 was phenotypically resistant to cefotaxime, ertapenem and amoxicillin/clavulanic acid (**Figure 4a**, **Supplementary Table 5**). *K. pneumoniae* M3 and M8 were phenotypically resistant to co-trimoxazole, but were not predicted to encode genes conferring resistance to the sulfonamide sulfamethoxazole or to trimethoprim (**Figure 4a**). M22 encoded only one trimethoprim-associated resistance gene, *dfrA12* (**Figure 4a**).

The beta-lactamase-encoding genes *bla*_OXA-1_ (16/31, 51.6%) and *bla*_LAP-2_ (1/31, 3.2%) were found among UCI KpSC isolates. Within all the Ugandan KpSC genomes, *bla*_OXA-1_ (46/137, 33.6%) predominated, followed by *bla*_SCO-1_ (5/137, 3.6%) and *bla*_LAP-2_ (4/137, 2.9%). The majority of our KpSC isolates were predicted to encode an acquired ESBL (27/31, 87.1%), with *bla*_CTX-M-15_ dominating (24/31, 77.4%); *bla*_CTX-M-14_, *bla*_CTX-M-3_ and a variant closely related to *bla*_CTX-M-15_ were found once each (1/31, 3.2%). All UCI ESBL-positive isolates were resistant to one or more beta-lactam antibiotics (**Figure 4a**). Across all the Ugandan KpSC genomes, 108/137 (78.8%) of all KpSC isolates were predicted to encode for at least one copy of an acquired ESBL: *bla*_CTX-M-15_ (104/137, 75.9%), *bla*_CTX-M-14_ (2/137, 1.5%), *bla*_CTX-M-3_ (1/137, 0.7%) and the variant of *bla*_CTX-M-15_ (1/137, 0.7%) (**Figure 4b**, **Supplementary Table 5**).

Few UCI isolates (5/31, 16.1%) and across the whole Ugandan KpSC genome dataset (7/137, 5.1%) were predicted to encode an acquired carbapenemase (**Figure 3a**, **Figure 4**). All carbapenemase-encoding isolates [*K. pneumoniae* M83 (NDM-7), M88 (NDM-5, OXA-232) and M99 (NDM-7); *K. quasipneumoniae* M91 (NDM-7)] were phenotypically resistant to imipenem, ertapenem and meropenem (**Figure 4a**). None of the Ugandan KpSC genomes was predicted to encode for carbapenemase plus acquired colistin resistance (**Figure 3a**), nor did they encode for acquired fosfomycin or tigecycline resistance (**Figure 4b**).

Two of the 31 (6.5%; M35 and M37) isolates that were phenotypically resistant to the second-generation fluoroquinolone ciprofloxacin were predicted to be sensitive to this antibiotic by Kleborate (0.25 mg/L; **Supplementary Table 5**). Isolate M35 was not predicted to encode acquired fluoroquinolone resistance genes, while M37 was predicted to encode an acquired *aac(6’)-Ib-cr.*v2 gene, harbouring W102R and D179Y mutations required for acetylation of fluoroquinolones (39).

### Klebsiella oxytoca

*K. oxytoca* M6, recovered from the paediatric ward, belonged to ST145, and was of unknown K and O locus types (best matches KL169 and OL15, respectively; Kleborate). Its genome was different from that of other KoSC bacteria reported for Uganda (31,32) (**Table 2**). M6 did not encode genes for colibactin, aerobactin, salmochelin or yersiniabactin. In line with its identification as *K. oxytoca*, M6 encoded the core class A beta-lactamase *bla*_OXY-2_, conferring resistance to cephalosporins (40). Phenotypically and genotypically, M6 was XDR (**Table 3**). It was predicted by antiSMASH to carry the *til* BGC, which encodes tilimycin (which in the presence of indole is converted to tilivalline) that contributes to antibiotic-associated haemorrhagic colitis (41); it is not known if isolate M6 produces tilimycin/tilivalline, but its genome encodes all 12 genes of the BGC, sharing 94.84% similarity with the reference BGC sequence from *K. grimontii* (**Supplementary Figure B**). M6 did not encode the *leup* BGC.

**Table 2.** Genomic data for Ugandan KoSC isolates.

| Isolate | Species | Source | Condition # | Accession | MLST \$ | *til* ^ | *leup* ^ | Reference |
| --- | --- | --- | --- | --- | --- | --- | --- | --- |
| M6 | *K. oxytoca* | Blood | BSI | SAMN53403839 | ST145 | + | - | This study |
| RSM7756 | *K. michiganensis* | Blood | BSI | GCA_030717465.1 | ST330 | - | + | (22) |
| RSM5061 | *K. michiganensis* | Blood | BSI | GCA_030490345.1 | ST748 | - | + | (22) |
| RSM9152 | *K. michiganensis* | Blood | BSI | GCA_030490325.1 | ST748 | - | + | (22) |
| RSM9152B-2 * | *K. michiganensis* | Blood | SM (BSI) | GCA_037202025.1 | ST748 | - | + | (23) |
| RSM6774 | *K. oxytoca* | Blood | BSI | GCA_030506035.1 | ST2 (CC2) | + | - | (22) |
| RSM5033 | *K. oxytoca* | Faeces | GI | GCA_030518375.2 | ST773 | + | - | (22) |
| RSM5034 | *K. oxytoca* | Faeces | GI | GCA_030544885.1 | ST773 | + | - | (22) |
| RSM6895 | *K. oxytoca* | Rectal swab | GI | GCA_030506015.1 | ST774 | + | - | (22) |

\* Metagenome-assembled genome.

**Table 3.** Summary of phenotypic and genotypic (AMRFinder) AMR data for *K. oxytoca* M6.

| Class | Subclass | Phenotypic resistance | Genotypic resistance * | Reference accession(s) |
| --- | --- | --- | --- | --- |
| Aminoglycoside | Gentamicin | Gentamicin | *aac(3)-IId* | WP_000557454.1 |
| Aminoglycoside | Streptomycin | Not tested | *aph(3'')-Ib* (52.06, 100), *aph(6)-Id* | WP_001082319.1, WP_000480968.1 |
| Aminoglycoside/<br>quinolone | Amikacin/kanamycin/<br>quinolone/tobramycin | Not tested | *aac(6')-Ib-cr* | WP_063840321.1 |
| beta-Lactam | beta-Lactam | Piperacillin/tazobactam,<br>amoxicillin/clavulanic acid | *blaTEM-1* | WP_000027057.1 |
| beta-Lactam | Carbapenem | Imipenem | *blaNDM-7* | WP_032495622.1 |
| beta-Lactam | Cephalosporin | Cefepime, cefotaxime,<br>ceftriaxone, ceftazidime | *blaOXA-1*, *bla<sub>CTX-M</sub>-15* | WP_001334766.1, WP_000239590.1 |
| Bleomycin | Bleomycin | Not tested | *ble* | WP_004201167.1 |
| Efflux | Efflux | Not tested | *emrD* (100, 94.16) | ACN65732.1 |
| Fosfomycin | Fosfomycin | Not tested | *fosA* (100, 88.49) | WP_004214174.1 |
| Phenicol | Chloramphenicol | Not tested | *catB3* (70, 100) | WP_000186237.1 |
| Quinolone | Quinolone | Ciprofloxacin | *oqxB* (100, 96.95),<br>*qnrB1* | WP_023323140.1, WP_014386481.1 |
| Sulfonamide | Sulfonamide | Co-trimoxazole | *sul2* | WP_001043260.1 |
| Tetracycline | Tetracycline | Not tested | *tetB* | WP_001089068.1 |

\* All shared 100% amino acid identity with the reference sequence with 100% reference coverage, unless shown otherwise (% coverage,% identity).

## DISCUSSION

The increasing prevalence of MDR *Enterobacterales*, combined with challenges in implementing effective infection control measures, poses significant concerns for cancer care in low-income settings. Our previous work (6) showed rates of antibiotic resistance were high among bacterial isolates recovered from hematologic cancer patients with febrile neutropenia at the UCI but, for example, potential transfer of MDR bacteria between patients could not be determined based solely on phenotypic data. In this study, WGS data were generated for 66 Gram-negative isolates (*E. coli* and *Klebsiella* spp.) previously recovered from the blood of cancer patients with febrile neutropenia (6).

Transportation of DNA samples from low-resource to high-resource settings for WGS is now commonplace, but publications describing such work typically report DNA extraction and sequencing protocols while omitting specific details about shipping. Bacterial DNA remains stable following ambient shipping of enriched cultures, isolates dried onto filter paper, and thermal lysates (“boilates”) (42,43). We show CTAB-extracted bacterial genomic DNA remained stable in nuclease-free water through 3 days of shipping from Uganda to the UK at ambient temperature. Although there were a small number of samples which arrived with very low volumes post-shipping due to evaporative losses, these could be prevented by the use of tightly closed screw top tubes and/or parafilm seals. This is a feasible and relatively low-cost option for researchers in low-resource settings to access affordable commercial, high-throughput WGS to characterise BSL-2 organisms (i.e. moderate-risk pathogens associated with human diseases that are usually treatable and not commonly spread via aerosols).

While cost-effective and used widely in low-resource settings, phenotypic testing has limitations including the potential for misidentification of closely related or rare bacteria (8,44). We noted discordance between conventional phenotypic and *rpoB*-gene-based identification of some of our isolates. This is a recognised but important under-reported issue. The range of phenotypic tests used in low-resource settings should be increased to improve species identification of emerging pathogens. For example, *K. oxytoca* isolates can be differentiated from *E. coli* based on the former being non-motile, urease-positive and citrate-positive (40). Species of the KpSC can only be distinguished using molecular methods or MALDI-TOF-MS (44,45). Wider adoption of multiplex PCR-based approaches for identification of KpSC isolates will improve our understanding of their routes of transmission and epidemiology in clinical settings (44).

In general, isolates were not similar enough to one another for us to infer transmission events within and between wards based on clinical outbreak-oriented relatedness thresholds (≤20 SNPs) (46,47), with >100 SNPs difference observed among many of our closely related isolates. *E. coli* M32 and M40, isolated 6 months apart from patients in the paediatric ward, differed by only 64 SNPs and shared identical AMR genotypes and phenotypes. The recent DRUM study used 0, 1, 2, 5 or 10 SNPs difference between ESBL *E. coli* genomes to demonstrate patient transmission (47). However, a recent One Health study proposed assessing genomic relationships of cross-source clusters and potential transmission events at ≤100 SNP threshold to detect linkages otherwise obscured by a ≤20 SNP threshold (46). This approach may well be applicable when assessing persistence of infection-associated isolates in the built environment of clinical wards.

STs detected in UCI *E. coli* genomes were broadly similar to those in the wider Ugandan dataset analysed here, with ST167 predominating and ST131 also well-represented. While 9.1% of the Ugandan genomes analysed were ST10, and ST10 dominated ESBL *E. coli* from Uganda in the DRUM study (8.5%) (47), ST10 was absent from the UCI *E. coli* genomes. The AMR prevalence and genotypes reported in this study are broadly similar to those from other Ugandan *E. coli* studies: MDR phenotypes with *bla*_CTX-M-15_ and *bla*_OXA-1_ are frequently seen, while carbapenem resistance remains rare and is typically linked to *bla*_NDM_ when present (23,25,26,48). The recent extensive DRUM study in Uganda and Malawi found *bla*_CTX-M-15_ in 68.4% and *bla*_CTX-M-27_ in 14.3% of community-derived ESBL *E. coli* isolates (47). Our WGS data from bacteraemia isolates showed similar trends: *bla*_CTX-M-15_ (61.7%) and *bla*_CTX-M-27_ (8.8%).

Phenotypic resistance to carbapenems was rare (6), and genes typically conferring carbapenem resistance were mostly absent from UCI *E. coli*. *bla*_NDM-5_-positive isolates (4/34; 11.76%) were phenotypically resistant to ertapenem, imipenem and meropenem. *bla*_NDM-7_ was present in isolate M64, also phenotypically carbapenem-resistant. Isolates M17 and M23 were phenotypically resistant to carbapenems in the absence of carbapenemase-encoding genes. The most likely explanation for this is hyperproduction of ESBLs (*bla*_CTX-M-55_ and *bla*_CTX-M-27_ were detected in isolates M17 and M23, respectively) combined with downregulation of porin genes, through mechanisms that are usually missed by Kleborate. ESBL-positive ST131 *E. coli* isolates from bloodstream infections (BSIs) have previously been shown to develop carbapenem resistance through these mechanisms (49), and BSI with carbapenemase-negative, carbapenem-resistant *E. coli* were associated with prior exposure to cefepime and ceftazidime-avibactam in cancer patients in the USA (50). *bla*_CTX-M-27_ is strongly associated with *E. coli* ST131, one of the most virulent and globally disseminated extraintestinal pathogenic *E. coli* (ExPEC) lineages (51). In this study, 5/34 *E. coli* isolates were ST131. Of the three isolates encoding *bla*_CTX-M-27_, two (M23 and M70) were ST131. The presence of *bla*_CTX-M-27_ defines the C1-M27 subclade of ST131 (52), which is globally expanding (53) but so far under-represented in reports from sub-Saharan Africa, likely due to limited access to the WGS that is necessary to differentiate *bla*_CTX-M-27_ from other ESBLs (53,54).

Carbapenem-resistance-conferring genotypes were also rare (3/386) among the publicly available Ugandan *E. coli* genome data analysed here, while cephalosporin resistance determinants, predominantly *bla*_OXA-1_, were more common (89/386). The relatively high prevalence of genotypes associated with gentamicin resistance [particularly *aac*(3)II genes] and ceftriaxone resistance (particularly *bla*_CTX-M-15_) in *E. coli* isolates is concerning, given the reliance on these together with penicillin-based drugs (cloxacillin and benzylpenicillin) as broad-spectrum first-line options for the treatment of septicaemia in the <u>Uganda Clinical</u> <u>Guidelines</u>. Among the UCI-derived *E. coli* isolates included in this study, 97% (33/34) had MDR or XDR phenotypes, defined here as resistance to at least one agent from three or four different classes of antibiotic, respectively. This highlights the importance of urgent blood cultures and antibiotic susceptibility testing when septicaemia is suspected in vulnerable cancer patients, to refine the treatment of specific cases while also collating valuable data for antibiograms to inform updates to local empiric prescribing guidelines.

While the WHO GLASS database provides some insights into antibiotic resistance rates in bacteriologically confirmed *E. coli* BSIs in the Africa region (e.g. meropenem 8.3%, ceftriaxone 74.0%, ciprofloxacin 63.8% – 2023 data, <u>the latest available at the time of</u> <u>writing</u>), few data from Ugandan health facilities have been submitted to this database. Only 21/47 (44.7 %) of bacteriologically confirmed BSIs submitted to WHO GLASS from Uganda in 2023 were attributed to *E. coli.* Among these, resistance to meropenem remained low (5.6%), while resistance to ceftriaxone and ciprofloxacin was high (81.2% and 91.7%, respectively).

The 26 different STs identified across 31 UCI KpSC isolates reflect the diversity in profiles seen country-wide (106 different STs; 329 available profiles from Uganda) and reported globally. A recent extensive analysis of global genomic diversity among 22,569 *K. pneumoniae* isolates from humans included 847 from Africa, but none from Uganda (55). The most prevalent STs (ST17; ST39; ST307), O types (O1ɑβ,2ɑ; O1ɑβ,2β) and K types (K62, K25, K2) reported from Africa (55) were well-represented in the Ugandan KpSC genomes analysed here.

The virulence factors predicted in our genomes are broadly similar to those reported for KpSC genomes from Uganda, with no *rmp*ADC hypermucoidity locus (also rare in the national dataset), no colibactin, 45.2% with yersiniabactin (46.0% in the national dataset) and low Kleborate virulence scores. Hypervirulent *Klebsiella pneumoniae* clones remain rare in Ugandan KpSC genomes, but some high-risk STs are represented and there is high carriage of genes conferring resistance to multiple classes of antibiotics (mean 7.3 antibiotic classes in our genomes and 6.5 classes in the national dataset). In resource-limited settings, preventing transmission and successfully treating infections caused by these organisms is likely to be challenging.

Almost all our KpSC isolates had ESBL-encoding and other resistance genes consistent with their MDR (19%) and XDR (74%) phenotypes. The phenotypic susceptibility of the majority of isolates to carbapenems was accurately predicted by Kleborate. Only four isolates had carbapenemase-encoding genes (*bla*_NDM-7_, n=3; *bla*_NDM-5_ with *bla*_OXA-232_,n=1); all four were phenotypically resistant. Efflux overexpression and porin loss could explain the phenotypic resistance to fluoroquinolones in M35, in the absence of the fluoroquinolone resistance genes Kleborate looks for (56).

Of the two *K. pneumoniae* isolates belonging to high-risk STs in our study, M15 (ST307), which carried *bla*_CTX-M-15_ and *bla*_OXA-1_, lacked carbapenemase-encoding genes and was susceptible to carbapenems as well as cefoxitin and piperacillin/tazobactam, while isolate M88 (ST147) had *bla*_NDM-5_, *bla*_OXA-232_ and other genes consistent with its XDR phenotype (resistant to all tested drugs). Although the virulence scores from Kleborate were low (0 or 1) for all of our and all but two of the Ugandan KpSC genomes, the presence of high-risk KpSC STs, and very high prevalence of MDR/XDR phenotypes driven by a range of mechanisms in isolates from this patient group, is a cause for serious concern and future management of BSIs in such patients should take these findings into consideration. In the UCI, and nationally, antibiograms should be developed and antibiotic-prescribing guidelines should be reviewed because a high proportion of KpSC genomes in this study and previous Ugandan studies encode genes conferring resistance to first- and second-line drugs for empiric treatment of BSIs, including gentamicin, penicillins, chloramphenicol and 3rd-generation cephalosporins (<u>Uganda Clinical Guidelines</u>). Moreover, ST307 has been shown to persist in the hospital environment, highlighting the critical need for infection prevention and control at the UCI (57). Given the genomic plasticity and global spread of KpSC, use of WGS surveillance with tools like Kleborate is important to spot emerging clones with both hypervirulence traits and genes conferring resistance to multiple antibiotic classes (16).

Herein we provide genome data for a clinical *K. oxytoca* isolate. The KoSC represents an emerging group of pathogens, notable for the bioactive molecules (tilimycin and tilivalline; leupeptin, pyrazinones and pyrazines) some strains produce that contribute to their virulence (18,40). Isolate M6 was not only XDR, but also predicted to encode a complete *til* BGC, as were other *K. oxytoca* isolates found in Uganda. This BGC – encoding for the virulence factor tilimycin – is common in *K. oxytoca*, *K. grimontii* and *K. pasteurii*, but rare in *K. michiganensis* isolates (19). The Ugandan *K. michiganensis* isolates encoded a complete *leup* BGC, in agreement with our recent large-scale analysis of the KoSC that demonstrated ∼90% of *K. michiganensis* genomes encode this genetic element (18). As of 20 November 2025, KoSC MLST data were sparse for African countries: Tunisia, ST220 (n=1 *K. michiganensis*); Ghana, ST419 (n=2 *K. oxytoca*); Nigeria, ST311 (n=1 *K. pasteurii*) and ST642 (n=1 *K. michiganensis*); Egypt, ST576 (n=5 *K. grimontii*), ST27 and ST542 (n=1 and n=4 *K. michiganensis*, respectively), and ST569 and ST629 (n=3 and n=1 *K. pasteurii*, respectively); Ethiopia, ST243 (n=1 *K. oxytoca*); United Republic of Tanzania, ST520 (n=1 *K. oxytoca*) and ST533 (n=1 *K. michiganensis*); South Africa, ST350 (n=1 *K. grimontii*), ST29, ST170, ST232 and ST242 (n=1, n=2, n=1 and n=1 *K. michiganensis*, respectively) and ST450 (n=3 *K. oxytoca*). Consequently, we have added all the MLST data we generated for Ugandan KoSC isolates to PubMLST.

Given the resource constraints in Uganda, with limited access to blood culture and antibiotic susceptibility testing for the majority of patients with suspected BSIs, national and facility-level data should be leveraged as far as possible to refine empiric treatment guidelines. Despite recognised limitations in using genomic data to predict AMR phenotypes (7,58), bacterial genomes derived from research studies will provide a valuable supplementary resource for informing policymakers about prevalent resistance determinants in circulation and for guiding the development of cost-effective and accessible rapid diagnostic tests targeting the most common genetic drivers of AMR.

## ACKNOWLEDGEMENTS

We thank the Institut Pasteur teams for the curation and maintenance of BIGSdb-Pasteur databases at https://bigsdb.pasteur.fr/.

## AUTHOR CONTRIBUTIONS

Conceptualization: ML, JW, LH; Data Curation: ML, JW, ALM, LH; Formal analysis: ML, JW, LH; Funding acquisition: ML, JW, LH, BA; Investigation: ML, JW, ALM, LH, DPK, FB, EK, LK, BA, GK, FL, SS, NN, JO, HD, JK, WP; Methodology: ML, JW, ALM, LH, DPK, FB, EK, LK, BA, GK, FL, SS, NN, JO, HD, JK, WP; Project administration: ML, JW, ALM, LH, DPK, FB, EK, LK, BA, GK, FL, SS, NN, JO, HD, JK, WP; Resources: ML, JW, ALM, LH, DPK, FB, EK, LK, BA, GK, FL, SS, NN, JO, HD, JK, WP; Software: ML, JW, LH; Supervision: JW, LH, DPK, FB, BA, JO, HD, JK, WP; Validation: ML, JW, LH; Visualization: LH; Writing- original draft: ML, JW, LH; Writing- reviewing and editing: ML, JW, ALM, LH, DPK, FB, EK, LK, BA, GK, FL, SS, NN, JO, HD, JK, WP

## Abbreviations

3GCRE: third-generation cephalosporin-resistant *Enterobacterales*
AMR: antimicrobial resistance
ANI: average nucleotide identity
BGC: biosynthetic gene cluster
BSI: bloodstream infection
CRE: carbapenem-resistant *Enterobacterales*
ESBL-PE: extended-spectrum-beta-lactamase-producing *Enterobacterales*
KoSC: *Klebsiella oxytoca* species complex
KpSC: *Klebsiella pneumoniae* species complex
MDR: multidrug-resistant
ST: sequence type
UCI: Uganda Cancer Institute
WGS: whole-genome sequencing
XDR: extensively drug-resistant

## Data availability statement

The sequence data reported on in this article are available under BioProject PRJNA1369824. Additional supplementary materials are available from https://figshare.com/projects/_b_Comparative_analyses_of_Gram-negative_bacteria_isolated_from_cancer_patients_with_bacteraemia_at_the_Uganda_Cancer_Institute_b_/272401. Data have also been deposited with Pathogenwatch: [*E. coli*](https://pathogen.watch/collections/rcyZbWvEqiuFRcM13MZY98-escherichia-coli-uci); [*K. oxytoca*](https://pathogen.watch/collections/nmNLJbgteKNjE5ozqpj1VB-klebsiella-oxytoca-uci); [*K. pneumoniae*](https://pathogen.watch/collections/cmXuaVsC6fHcPgtMYGPWvQ-klebsiella-pneumoniae-uci); [*K. variicola*](https://pathogen.watch/collections/oqgQRPUNU9tfrt9PzKsbJs-klebsiella-variicola-uci); [*K. quasipneumoniae*](https://pathogen.watch/collections/aV8mbcoUBzLPpT51GGCRYB-klebsiella-quasipneumoniae-uci).

---

**Supplementary Figure A.**
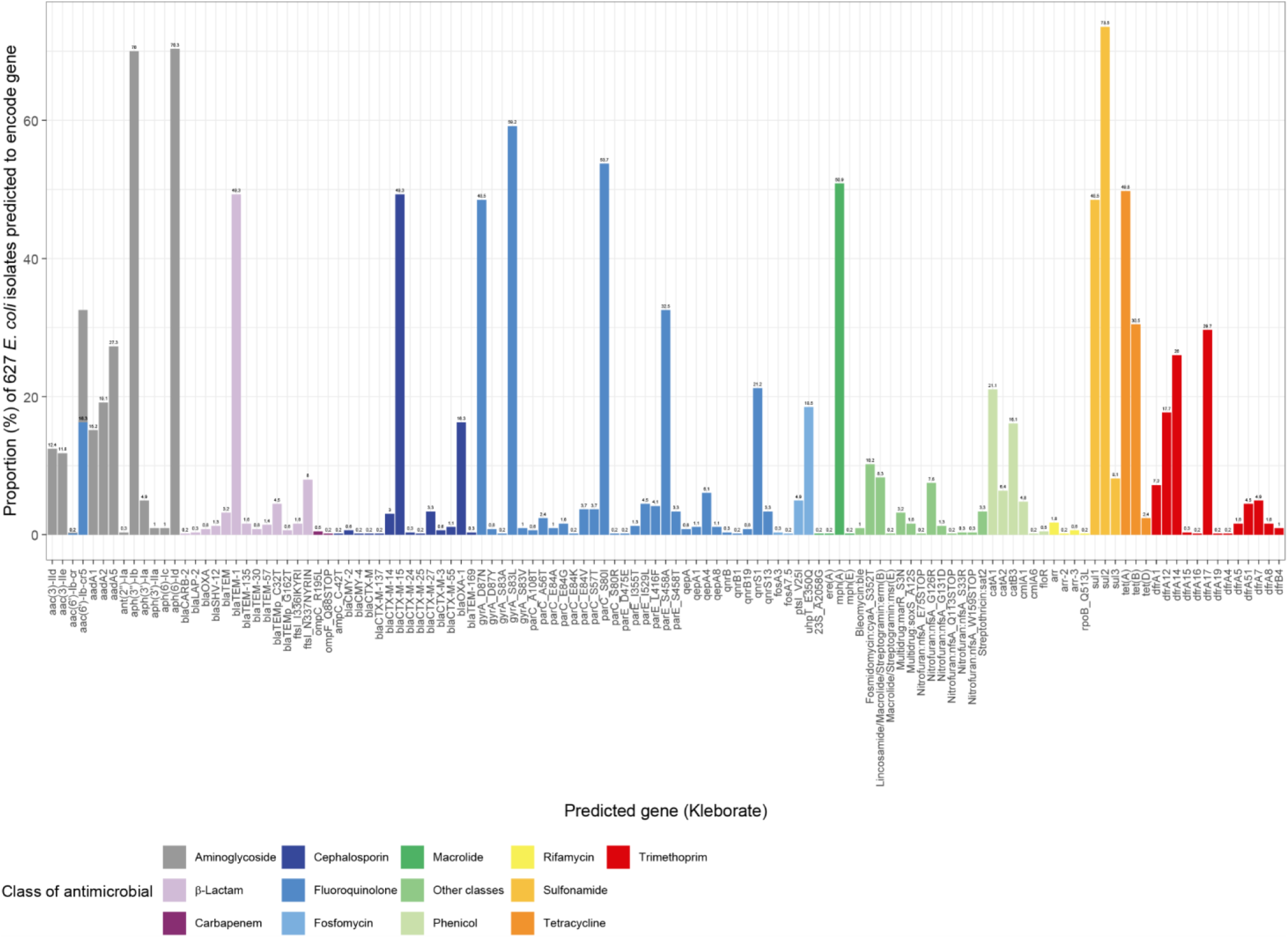
Summary of AMR gene predictions generated by Kleborate for the 627 *E. coli* publicly available genomes from Uganda.

**Supplementary Figure B.**
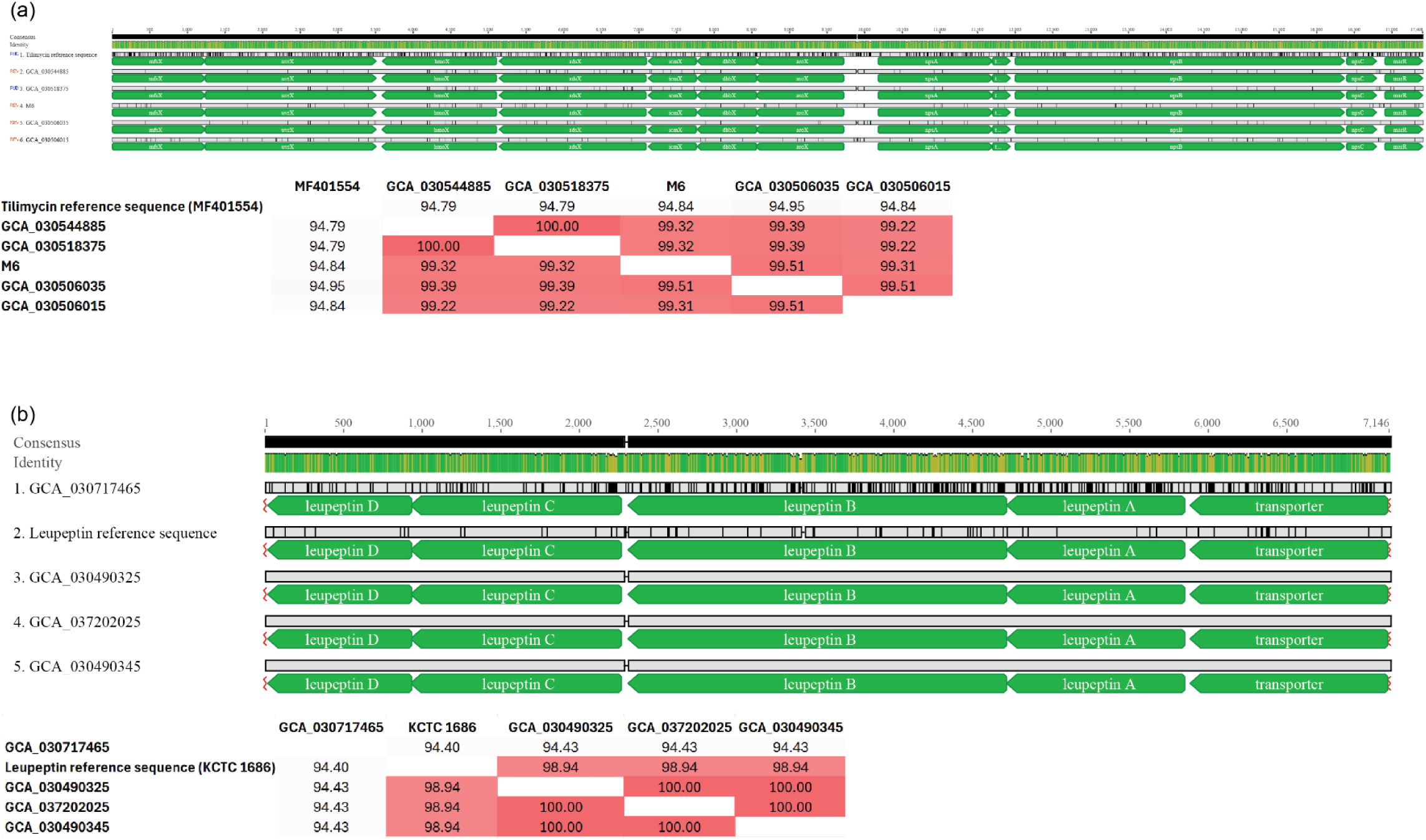
Confirmation that complete *til* and *leup* BGCs are encoded by members of the KoSC isolated in Uganda. (a) Detection of the complete *til* BGC in *K. oxytoca* genomes, with nucleotide similarity (%) matrix. (b) Detection of the complete *leup* BGC in *K. michiganensis* genomes, with nucleotide similarity (%) matrix. Multiple-sequence alignments were generated using Clustal Omega 1.2.2 and visualized using Geneious Prime, and are available to download from figshare along with antiSMASH outputs (doi:10.6084/m9.figshare.31546765, doi:10.6084/m9.figshare.31545100, doi:10.6084/m9.figshare.31541713).

